# Quantification of Rho-termination *in vivo* using qRT-PCR: a comprehensive analysis

**DOI:** 10.1101/2023.04.11.536429

**Authors:** Monford Paul Abishek N, Heung Jin Jeon, Xun Wang, Heon M. Lim

## Abstract

In prokaryotes, the Rho protein mediates Rho-dependent termination (RDT) by identifying a non-specific cytosine-rich Rho utilization site on the newly synthesized RNA. As a result of RDT, downstream RNA transcription is reduced. Due to the bias in reverse transcription and PCR amplification, we were unable to identify the RDT site by directly measuring the amount of mRNA upstream and downstream of RDT sites. To overcome this difficulty, we employed a 77 bp reporter gene *argX*, coding transfer RNA that binds L-arginine, tRNA^arg^ from *Brevibacterium albidum*, and transcriptionally fused it to the sequences to be assayed. We constructed a series of plasmids by combining a segment of the galactose (*gal*) operon sequences, both with and without the RDT regions at the ends of cistrons (*galE*, *galT,* and *galM*) upstream of *argX*. The RNA polymerase will transcribe the *gal* operon sequence and *argX* unless it encounters the RDT encoded by the inserted sequence. We observed similar tRNA^arg^ half-lives expressed in these transcriptional fusion plasmids. Therefore, the amount of tRNA^arg^ directly represents the number of transcripts transcribed. Using this approach, we were able to effectively assay the RDTs in the *gal* operon by quantifying the relative amount of tRNA^arg^ using quantitative real-time PCR (qRT-PCR) analyses. The resultant RDT% for *galET*, g*alTK*, and at the end of *galM* were 36, 26, and 63, individually. Our findings demonstrate that combining tRNA^arg^, with qRT-PCR can directly measure RDT efficiencies *in vivo*, making it a useful tool for gene expression research.

## IMPORTANCE

Rho is a protein that is crucial for bacterial transcription termination. It plays an important role in the release of mRNA by binding to target sites on nascent RNA and causing transcription termination. The difficult molecular mechanisms and real-time nature of transcription termination make studying Rho-dependent termination (RDT) *in vivo* hard. Here, we establish and validate the significance and utility of using tRNA^arg^ as a reporter gene to detect Rho-dependent termination (RDT) *in vivo*. This method, which looks at RNA expression, reflects only the transcription frequency of the gene to which it is fused, is more direct than employing β-galactosidase or fluorescent proteins as reporter genes, and should be promoted and widely used to detect transcription termination. Thus, this method offers a reliable and quantitative method for examining the role of Rho and other factors involved in transcription termination.

Two forms of transcription termination procedures have been reported in bacteria (1, 2): 1) Intrinsic termination or Rho-independent termination (RIT) by the terminator hairpin formed on the transcript at the end of the last transcriptional unit and 2) Rho-dependent termination (RDT) (reviewed in (2)). The Rho protein binds to a specific sequence in the mRNA transcript and moves along the RNA chain to the 3’ end, where it causes the RNA polymerase (RNAP) to dissociate from the template DNA strand, causing transcription to stop (3). In *Escherichia coli*, Rho is essential for survival, and about 20-30% of its genes are terminated by Rho, indicating its importance in *E. coli* (4, 5). Although researchers have been studying the working mechanisms of Rho for decades through genetic and biochemical experiments, several issues make predicting and determining RDT regions *in vivo* still difficult (3, 6, 7), including (i) mechanism complexity of RDT involves multiple components, including RNAP, the nascent RNA transcript, and the Rho protein. The interactions between these components make understanding the mechanism of RDT difficult; (ii) Rho is not specific to any mRNA sequence and can bind to any RNA sequence containing a Rho utilization (*rut*) site; (iii) the instability of Rho-terminated RNA 3’ ends: the 3’ end of Rho-terminated mRNA is rapidly digested by 3’ to 5’ exonuclease digestion after RDT due to the lack of secondary structure.

Our previous research recognized three RDTs that span the *galE-galT* and *galT-galK* cistron junctions, the first and second structural genes of the galactose operon, as well as downstream of *galM*, the last structural gene of the *gal* operon (1, 8-10). Using synthetic small RNAs (sysRNAs), we were also able to identify the *in vivo* RDT site downstream of *galM* (6), however, we were unable to identify the RDT sites covering the *galE-galT* and *galT-galK* cistron junctions, possibly due to ribosome and sysRNA competition. Therefore, to gain a deeper insight into the RDT mechanism, it is necessary to carefully consider these issues and apply more widely applicable techniques for RDT identification.

The quantitative real-time PCR (qRT-PCR) technique is used to determine the amount of mRNA in a sample, which can be used as an indicator of gene expression (11, 12). qRT-PCR is widely used in many research areas, including gene expression profiling, validation of gene expression arrays, disease diagnosis, drug development, and forensic analysis (13-15). qRT-PCR was used to validate RDT sites. Chhakchhuak and Sen designed multiple qRT-PCR primer probes and compared the gene expressions of Rho function-deficient strains relative to the wild-type strains, and the upregulation of the genes implied the presence of RDTs upstream of these probe sites (16). Notably, this approach does not reflect changes in the relative amount of transcripts upstream and downstream of RDT sites; it reflects changes in the relative amounts of gene expression between the wild type and Rho function-deficient strains. Due to bias in reverse transcription and PCR amplification, the quantities between two different transcripts cannot be directly compared (17, 18). Therefore, researchers cannot directly compare the difference in transcript amounts between the upstream and downstream RDT sites.

We wondered if there is a way to fairly show the amount of transcription between the upstream and downstream of RDT sites, which would more directly show transcription termination and could also be more easily used to calculate transcription termination efficiency. In this study, we employed a transcriptional reporter system, tRNA^arg^ from *Brevibacterium albidum*, which was transcriptionally fused to the sequences to be assayed. With this approach, transcription initiated from the *gal* operon promoters transcribes the inserted *gal* sequence, tRNA^arg^, unless it is terminated by an upstream transcription termination that is aimed to be assayed. We assayed the RDT region by quantifying the relative amount of tRNA^arg^ using qRT-PCR analyses. With this approach, we observed a gradual decrease in *gal* operon transcripts due to RDT and we were also able to calculate the efficiency of RDT. This approach represents a valuable tool for advancing future investigations into RDT and gene expression.

## RESULTS

### Establishment of RDT in vivo detection system

We would like to assay RDTs in *gal* operon by qRT-PCR using a pBR322-derived plasmid harboring a 77 bp gene *argX* coding for tRNA^arg^ from *B. albidum* (Fig. 1, Fig. S1). We chose tRNA^arg^ reporters from *B. albidum* because it was previously shown to be orthogonal to tRNAs in *E. coli* and was quite stable and could be detected by qRT-PCR analysis (19, 20). First, we tested the orthogonality of tRNA^arg^. Indeed, tRNA^arg^ was not detectable in MG1655-pKK232 where the *argX* gene was absent (data not shown), proving that tRNA^arg^ is orthologous to tRNAs in *E. coli*. Then, the half-life of tRNA^arg^ in the MG1655-pHL1141 strain was measured by qRT-PCR. Contrary to our expectation, the tRNA^arg^ did not show strong stability, instead, it had a similar half-life to *gal* mRNA, about 1.2 min (Table 1). We also tested the half-life of *cat* mRNA, as well as the half-life of 16S rRNA, and the results showed that the half-life of *cat* mRNA was about 1.2 min and 16S rRNA was very stable, with no degradation observed (Table 1).

**Figure 1.**
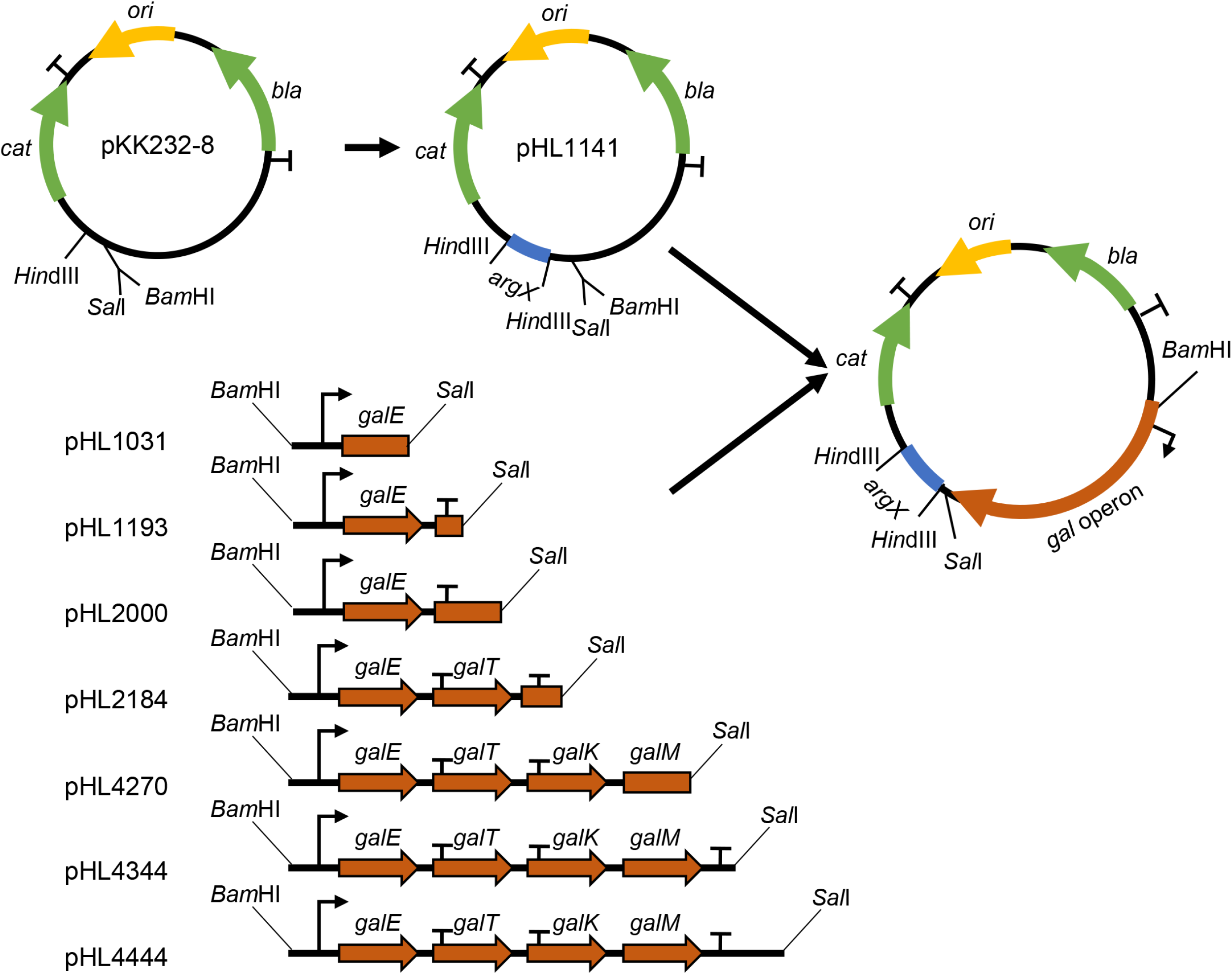
Plasmids used to measure RDT efficiency *in vivo* and their construction strategy. The plasmid map of pKK232-8; *ori*, plasmid origin of replication (yellow); Genes: *cat* and *bla*, Chloramphenicol acetyltransferase and β-lactamase (green) antibiotic resistance genes; *rrnBT1/T2*, transcription terminators are denoted as a (T); the restriction enzymes *Hin*dIII, *Sal*I, and *Bam*HI are indicated. Sequences of *argX* corresponding to tRNA^arg^ inserted upstream of the *cat* gene are shown in blue to generate pHL1141. Schematic presentation of the DNA fragments of *gal* operon harboring from -73 to 1,031, -73 to 1,193, -73 to 2,000, -73 to 2,184, -73 to 4,270, -73 to 4,344, and -73 to 4,444 were cloned in front of the tRNA^arg^ gene of pHL1141. The RDT site in the *gal* DNA fragments is represented by (T) at the end of the *gal* cistrons. The relative amounts of tRNA^arg^ from cells harboring pHL1031 (before-RDT), pHL2000 (before-RDT), pHL4270 (before-RIT/RDT), pHL1193 (after-RDT), pHL2184 (after-RDT), pHL4,344 (after-RIT), and pHL4,444 (after-RIT/RDT) were taken to represent the efficiency of RIT and RDT.

**Table 1.** Various RNA half-lives.

| RNA name | half-life (min) |
| --- | --- |
| tRNA<sup>arg</sup> | 1.2±0.09 |
| *galE* | 1.1±0.17 |
| cat | 1.5±0.14 |
| 16S rRNA | Stable (>4000) |

We then fused *argX* downstream of various *gal* operon sequences to construct seven pHL1141-derived plasmids (Fig. 1) and transformed them in *E. coli* MG1655 strains as described in Materials and Methods. We tested the half-lives of tRNA^arg^ expressed in these strains. The results showed that tRNA^arg^ half-lives were similar, ranging from 1.0 to 1.3 min (Table 2). The amount of RNA that is quantified by qRT-PCR depends on both the rate of RNA synthesis and the rate of RNA degradation. Since tRNA^arg^ is degraded at a similar rate on these plasmids, we believe that the amount tRNA^arg^ from these strains would represent the amount of *gal* transcript initiated from the *gal* promoters before and after the putative RDT site, respectively.

**Table 2.** tRNA^arg^ half-lives.

| RNA name | From plasmid | half-life (min) |
| --- | --- | --- |
| tRNA<sup>arg</sup> | pHL1031 | 1.2±0.10 |
|  | pHL1193 | 1.0±0.08 |
|  | pHL2000 | 1.3±0.11 |
|  | pHL2184 | 1.3±0.09 |
|  | pHL4270 | 1.3±0.17 |
|  | pHL4344 | 1.2±0.12 |
|  | pHL4444 | 1.3±0.08 |

### RDT at the galE-galT cistron junction

We measured the efficiency of RDT by measuring the amount of the *gal* transcript RNA before and after the RDT event at 1,183 *in vivo* (10). To measure the transcription ’before-RDT,’ the portion of the *gal* operon was from -73 to 1,031. To measure the transcription ’after-RDT,’ the portion was from -73 to 1,193. The resultant plasmids were pHL1031 and pHL1193, respectively (Fig. 1). These regions of *gal* contain all *cis-*acting elements necessary for *gal* expression. Amounts tRNA^arg^ from pHL1031 and pHL1193 would represent the amounts of *gal* transcript initiated from the *gal* promoters before and after the RDT at 1,183, respectively. We measured tRNA^arg^ expression in these two strains harboring pHL1031 and pHL1193 plasmids individually. The results indicated that in strain MG1655-pHL1193, tRNA^arg^ was 64 ± 19% of the level found in MG1655-pHL1031 (Fig. 2A). This suggests that transcription termination occurs approximately 36% of the time at 1,183 due to Rho in MG1655 (Table 3). In HME60 (*rho*::Amp^R^) cells, where Rho is non-functional (8), the tRNA^arg^ expression level was almost the same as that in MG1655-pHL1031 (Fig. 2B), which confirms that the measurement was of RDT at the *galE*-*galT* junction.

**Figure 2.**
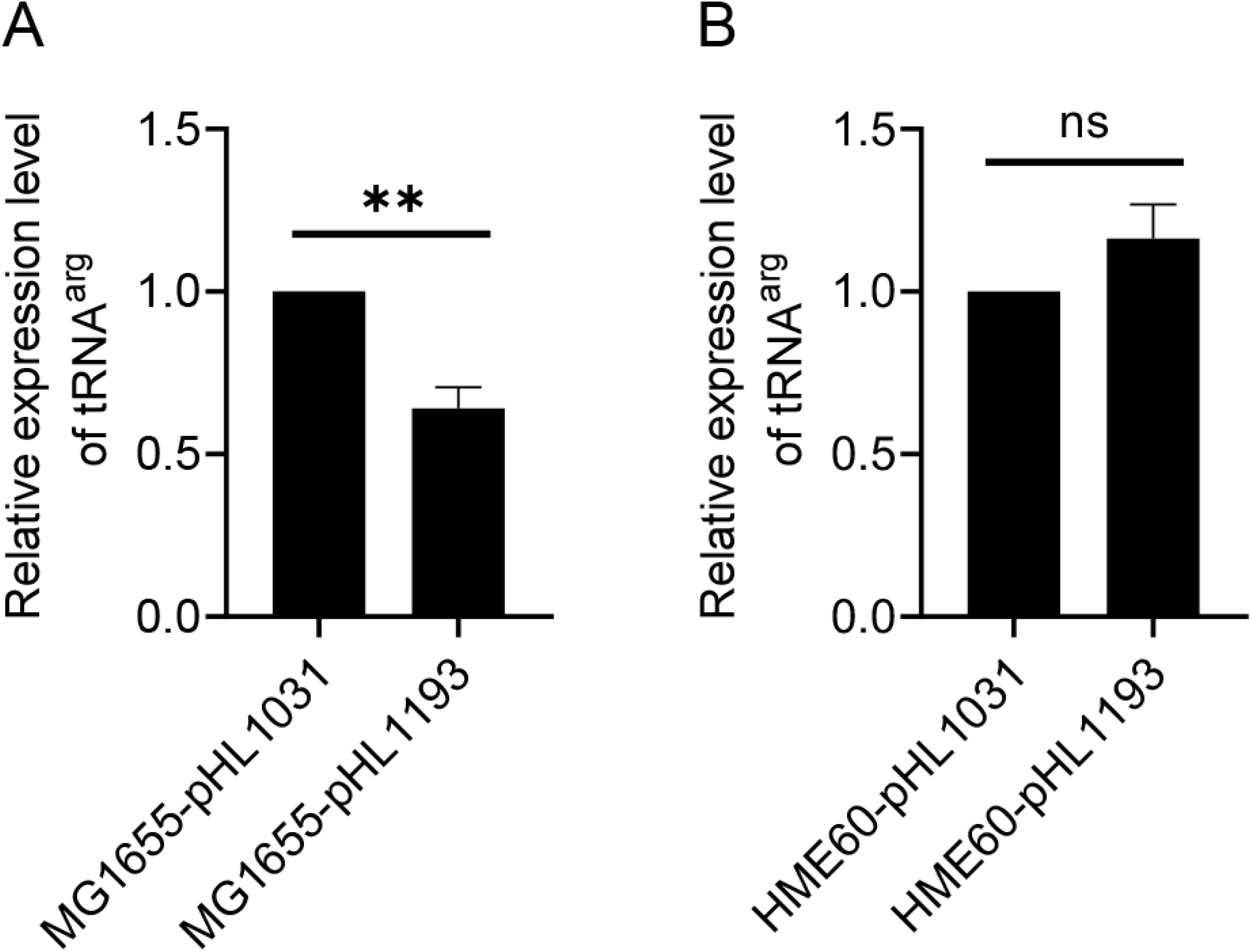
RDT at the *galE-galT* cistron junction. The results of qRT-PCR measurement of tRNA^arg^ in (A) MG1655 and (B) HME60 (*rho::*Amp^R^) are presented in which the relative amount of tRNA^arg^ in pHL1193 (after-*RDT*) is compared to that in pHL1031 (before-*RDT*). Error bars represent the mean fold-change ± standard deviation of three replicates from three independent experiments (n=3). ns: not significant, P value >0.05; **: significant differences, 0.001<P value < 0.01.

**Table 3.**
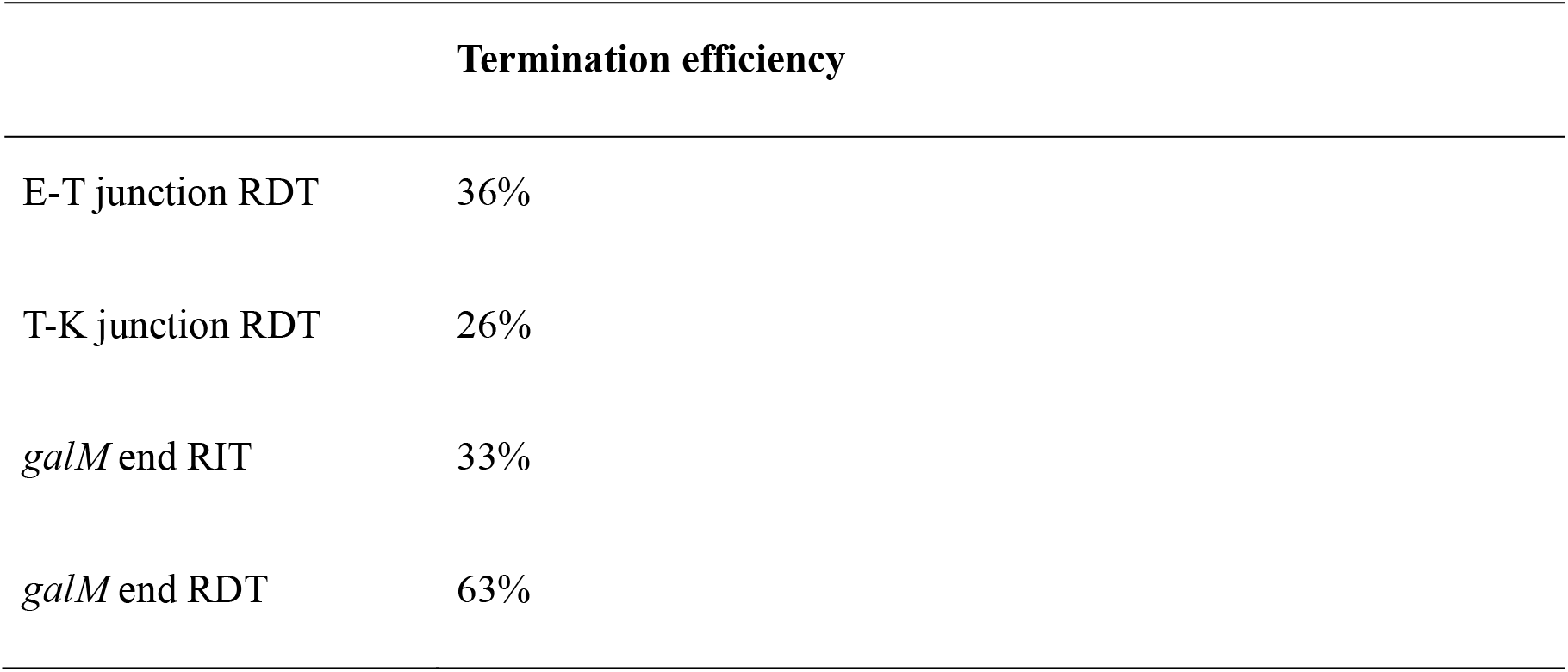
Termination efficiency calculated. The termination efficiency % was calculated using the formula: Termination frequency = 1 -read-through/upstream transcripts.

### RDT at the galT-galK cistron junction and Spot 42 enhanced it

We measured the efficiency of RDT that results in the generation of the mRNA *galET-short* at *2,121-2,125* (9). To measure transcription, a portion of the *gal* operon from -73 to 2,000 was cloned to generate the plasmid pHL2000. Another portion from -73 to 2,184 was cloned to generate the plasmid pHL2184 (Fig. 1). Since the cell-free assay demonstrated that Rho terminates transcription at 2,184 (9, 21), we anticipated that the amount of tRNA^arg^ would be less from MG1655-pHL2184 than MG1655-pHL2000. The result, indeed, showed that tRNA^arg^ in MG1655-pHL2184 is 74 ± 19 % of that in MG1655-pHL2000 (Fig. 3A), suggesting that Rho terminates transcription about 26 % of the time in the presence of the RDT site.

**Figure 3.**
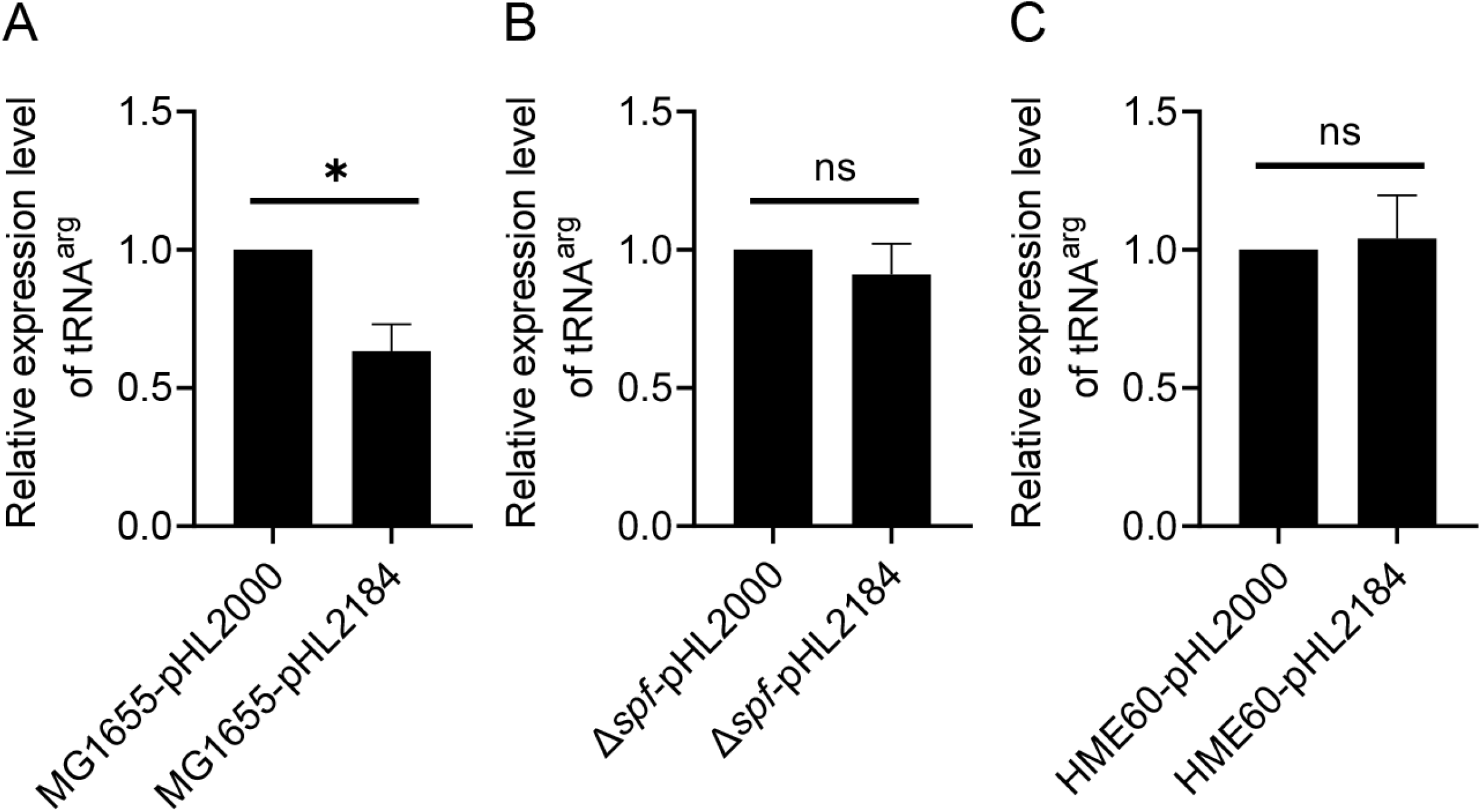
Spot 42 controls RDT at the *galT-galK* cistron junction. The results of qRT-PCR measurement of tRNA^arg^ in (A) MG1655, (B) MG1655Δ*spf*, and (C) HME60 (*rho::*Amp^R^) are presented in which the relative amount of tRNA^arg^ in pHL2184 (after-*RDT*) is compared to that in pHL2000 (before-*RDT*). Error bars represent the mean fold-change ± standard deviation of three replicates from three independent experiments (n=3). ns: not significant, P value >0.05; *: significant differences, 0.01<P value < 0.05.

Spot 42, a 109 nucleotide-long noncoding sRNA, binds to the middle of multi-cistronic mRNA at the *galT*-*galK* cistron junction and enhances RDT (22). Our qRT-PCR results showed that in the Δ*spf*-pHL2184 strain in which the Spot 42 coding gene was deleted, tRNA^arg^ after the RDT is 92 ± 10 % of that of Δ*spf*-pHL2000 (Fig. 3B), suggesting that in the absence of Spot 42, Rho hardly terminates transcription even when the RDT site is present. These results demonstrate that Spot 42 is the critical factor that causes RDT to generate the *galET* mRNA. We also observed that the tRNA^arg^ in HME60-pHL2184 and HME60-pHL2000 were almost equal, regardless of the presence of RDT sequences (Fig. 3C). The above findings with various strains demonstrated that measuring RDT efficiency with the qRT-PCR technique is quite effective.

### RDT and RIT at the end of the operon

At the end of *the gal* operon, transcriptions go through two terminators: the upstream RIT and downstream RDT (1). Rho terminates transcripts at 4,409 and has been quickly processed by exonuclease digestion (1). For measurement of transcription ’before-*RIT*-*RDT*’ (from -73 to 4,270), and transcription ’after-*RIT’* and ’after-*RIT*-*RDT’* (-73 to 4,344 and -73 to 4,444). The resultant plasmids were pHL4270, pHL4344, and pHL4444, respectively (Fig. 1). Since the half-lives of tRNA^arg^ expressed by MG1655-pHL4270, MG1655-pHL4344, and MG1655-pHL4444 strains were similar, about 1.3, 1.2 and 1.3 min, respectively (Table 1). Thus, the amount tRNA^arg^ from MG1655-pHL4270, MG1655-pHL4344, and MG1655-pHL4444 would represent the amount of *gal* transcript initiated from the *gal* promoters before and after the RIT and RDT sites, respectively. The RIT and RDT site is at 4,315 and 4,409, we expected the amount of tRNA^arg^ would be less from the plasmid containing the RIT and RDT site than from the plasmid without it.

The amount of tRNA^arg^ was measured using qRT-PCR in MG1655 cells harboring the plasmids. The results showed that tRNA^arg^ in MG1655-pHL4344 and MG1655-pHL4444 is 67 (± 2) and 25 (± 8) % of that in MG1655-pHL4270 (Fig. 4A), suggesting that RIT efficiency is 33 %, RDT efficiency is 63 %, and transcription terminated by both RIT and RDT is 75% (Table 3). In HME60 (*rho*::Amp^R^) cells, the results showed that the tRNA^arg^ in HME60-pHL4344 and HME60-pHL4444 is 68 (±) 6 and 90 (±) 14 % of that in HME60-pHL4270 (Fig. 4B). This suggests that RIT efficiency is 32 %, almost same as in MG1655-pHL4333 (Fig. 4A) and transcription terminated by both RIT and RDT drops down to 10 % (Table 3). The expression level of tRNA^arg^ in HME60-pHL4444 was nearly equal to that in HME60-pHL4270, suggesting that the measurement was of RDT at the end of the operon (Fig. 4B).

**Figure 4.**
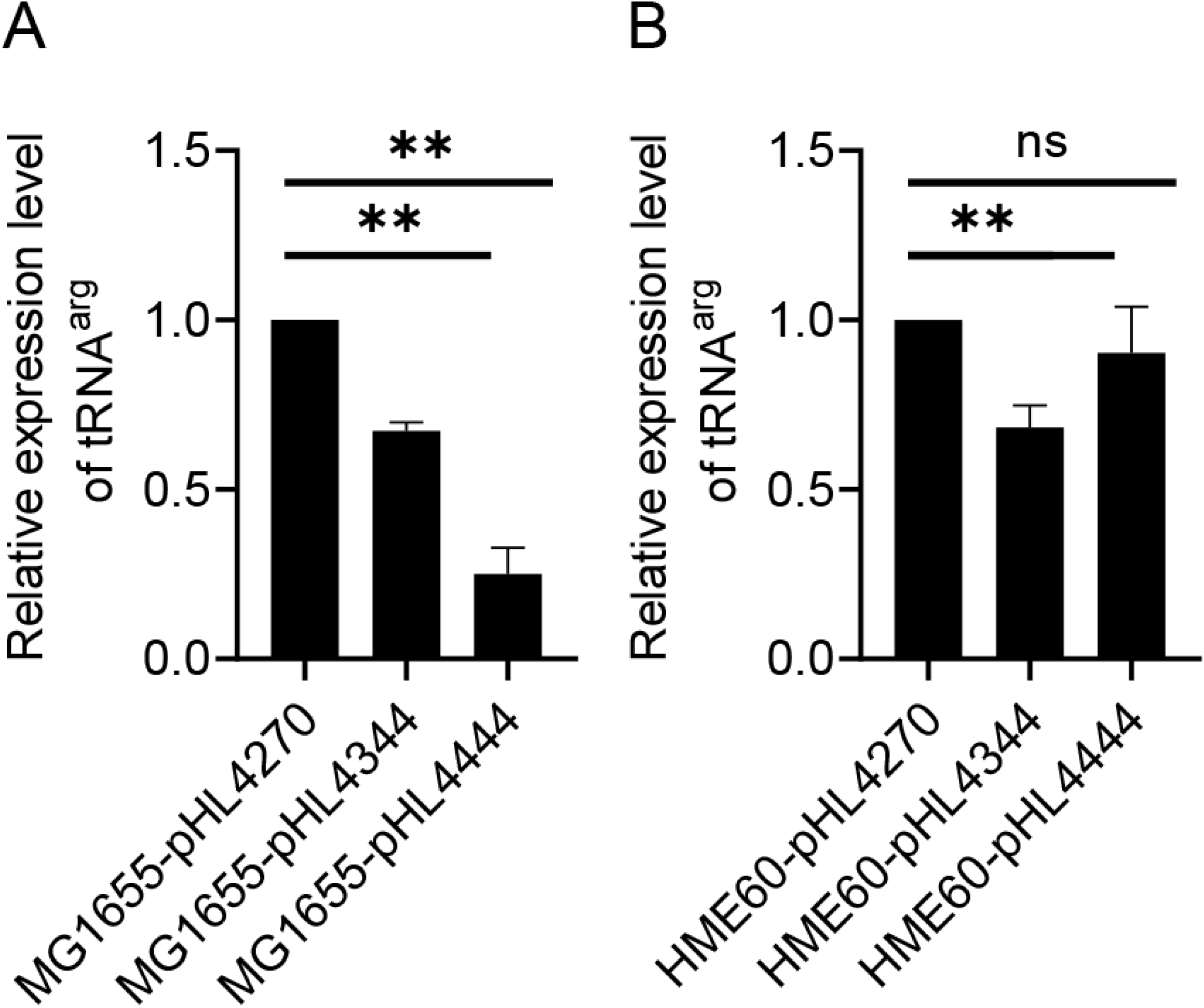
RDT at the end of *gal* operon. The results of qRT-PCR measurement of tRNA^arg^ in (A) MG1655 and (B) HME60 (*rho::*Amp^R^) are presented in which the relative amount of tRNA^arg^ in pHL4344 (after-*RIT*)/pHL4444 (after-*RDT*) is compared to that in pHL4270 (before-*RIT/RDT*). Error bars represent the mean fold-change ± standard deviation of three replicates from three independent experiments (n=3). ns: not significant, P value >0.05; **: significant differences, 0.001<P value < 0.01.

## DISCUSSION

In the present work, our results show that qRT-PCR analysis of tRNA^arg^ can directly reflect the amount of transcript before and after the RDT site, respectively. If RDT is effective, mRNA transcript levels should decrease after termination, indicating that the Rho protein has caused transcription termination (16, 23). Additional controls can be included, such as measuring transcript levels in cells that do not express Rho protein (here, HME60 (*rho*::AmpR)) or have mutations in the Rho-binding site. Inclusive, qRT-PCR can be a powerful tool for studying the role of the Rho protein in prokaryotic transcription termination, providing a quantitative measure of mRNA transcript levels before and after RDT (16, 23).

Transfer RNAs (tRNAs) are produced as precursors and then go through a multi-step maturation process to become mature tRNAs with a cloverleaf-shaped secondary structure (24, 25). The tRNAs are seasonably stable due to their unique secondary structure and the protective effect of the proteins bound to them (26). As a result, in prior studies, they were utilized as stable RNAs for reporter genes. However, in this research, we found that 77 bp tRNA^arg^ was not as stable as expected. This could be because tRNA^arg^ is a segment of a long transcript rather than a processed cloverleaf-shaped structure. As a result, it loses its initial secondary structure and physiological function, and it no longer has the reported stability.

Using tRNA^arg^ as a reporter gene to measure transcription has various advantages: (i) First and foremost, it avoids the bias inherent in translation. In gene expression studies, the RDT to be determined is typically fused with a gene that produces a protein, such as β-galactosidase or fluorescent proteins, and the intensity of the proteins serves as a measure of the transcriptional activity (27, 28). However, it has been shown that due to variations in translation rates, the quantity of transcription does not always correlate linearly with the amount of translation (29-31). Our findings show that tRNA^arg^ has similar half-lives across different plasmids, which explains why the tRNA^arg^ reporter precisely matches the rate of gene transcription. (ii) To quantify gene expression, the well-known β-galactosidase (*lacZ*) technique requires the addition of a substrate (such as X-gal), which can introduce unpredictability and be toxic to some cell types (32). In contrast, qRT-PCR assessment of tRNA^arg^ can detect smaller changes in gene expression levels and does not require an exogenous substrate, allowing for accurate and precise quantification of the effects of Rho-termination on gene expression.

In conclusion, qRT-PCR is a powerful tool for studying RDT in prokaryotes. Still, researchers should be aware of its limitations and select the best method suited to their experimental design and research question. Traditional methods for studying RDT, such as Northern blotting and primer extension assays, may be less sensitive, quantitative, and time-consuming (1, 6, 8, 10, 21, 22, 33). These methods may also necessitate more RNA and are unsuitable for high-throughput analysis. qRT-PCR is thus a powerful and efficient technique for studying RDT in prokaryotes, and its benefits make it an appealing option for researchers in this field.

## MATERIALS AND METHODS

### Bacterial strains, media, and growth conditions

For RDT analysis, *E. coli* MG1655, HME60 (*rho-15*), and MG1655 Δ*spf* strains were used. Chromosomal deletion strains of *E. coli* MG1655 were generated using phage Lambda Red-mediated recombineering (34). For plasmid construction, the DH5α strain was used. Primers used in this study are listed in Table S1. All cells were grown at 37°C in Lysogeny Broth supplemented with 0.5% (w/v) galactose and either chloramphenicol (15 μg/ml) or ampicillin (100 μg/ml) to reach an exponential growth phase with an optical density at 600 nm (OD_600_) of 0.6. Following that, RNA isolation was performed.

### Design and construction of plasmids

Sequences of *argX* from *B. albidum* corresponding to tRNA^arg^ were amplified by PCR using primer sequences shown in Table S1. The PCR fragment was digested with *Hin*d III, and then inserted immediately upstream of a promoter-less chloramphenicol resistance gene (*cat*) gene in pBR322 origin a medium-copy-number plasmid pKK232-8 (GE Healthcare, USA) to create the pHL1141 plasmid (8) (Fig. 1, Fig. S1) (20). To assay RDT, the *gal* operon DNA sequence was cloned between the *Bam*HI and *Sal*I restriction enzyme sites upstream of *argX* in pHL1141.

To generate the plasmids pHL1031, pHL1193, pHL2000, pHL2184, pHL4270, pHL4344, and pHL4444, the *gal* operon DNA sequences from *gal* coordinates -73 to 1,031, -73 to 1,193, -73 to 2,000, -73 to 2,184, -73 to 4,270, -73 to 4,333 and -73 to 4,444 were obtained with the corresponding primer pairs via PCR amplification (Table S1). The resulting PCR fragments were ligated to pHL1141, which has been digested with *Bam*HI and *Sal*I (Fig. 1). DNA sequencing was used to confirm the plasmid constructs.

### RNA preparation and Reverse transcription-quantitative PCR (qRT-PCR)

Following the manufacturer’s instructions, the clarified cell lysates (2×10^8^ cells) were used to purify total RNA using the Direct-zol RNA MiniPrep kit (Zymo Research, USA). One microgram of total RNA after Turbo DNase I (Thermo Fisher Scientific, USA) reaction to remove genomic or plasmid DNA was reverse-transcribed in a 20 μl reaction volume as previously described (8). The results from each sample were normalized using the internal control *rrsB*, which encodes 16S rRNA. The qRT-PCR reactions were set up in triplicates, and the mean Ct value was used to analyze three independent experiments further. The expression of each sample is presented as mean ±SD (35-37). The ΔΔCT values were subjected to a one-way analysis of variance (ANOVA) using the Bonferroni test. GraphPad Prism 9.0 was used for one-way ANOVA and graph plotting.

### RNA stability assay

We examined the decay rate of *galE*, tRNA^arg^, 16S rRNA, and *cat* mRNA species using wild-type (WT) MG1655 cells harboring plasmids (8) (as detailed in Tables 1 and 2). To determine RNA stability, we added 100 μg/mL of rifampicin, a transcription inhibitor (38), to LB-grown cells with an OD_600_ of 0.6. Total RNA was extracted for qRT-PCR analysis after samples were obtained at 0, 2, 4, and 8 minutes after rifampicin addition. We used 16S rRNA, to standardize the findings. The relative expression was calculated using the 2−ΔΔCT method by averaging the fold changes of three replicates from three independent experiments.

## SUPPLEMENTAL MATERIAL

Supplemental material is available online only (*PDF file*).

## ACKNOWLEDGEMENTS

This work was supported by the National Key Research and Development Program of China (2022YFF1000700), the National Natural Science Foundation of China (31971339 and 32171422), and the Fundamental Research Funds for the Central Universities (2662022SKYJ004). This work was also supported by the research fund of Chungnam National University. We declare no competing interests.

X.W., and H.M.L., Conceptualization, Formal analysis, Resources, Data curation, Funding acquisition, and Project administration; M.P.A.N., Investigation, Methodology, and Data curation; M.P.A.N., H.J.J., and X.W., Visualization, Validation, and Data analysis; M.P.A.N., X.W., and H.M.L., Writing - original draft, and Writing - review & editing.

